# Comparison of SARS-CoV-2 Indirect and Direct Detection Methods

**DOI:** 10.1101/2020.05.12.092387

**Authors:** Joel D. Pearson, Daniel Trcka, Sharon J. Hyduk, Marie-Ming Aynaud, J. Javier Hernández, Filippos Peidis, Suying Lu, Kin Chan, Jim Woodgett, Tony Mazzulli, Liliana Attisano, Laurence Pelletier, Myron I. Cybulsky, Jeffrey L. Wrana, Rod Bremner

## Abstract

The COVID-19 pandemic caused by the SARS-CoV-2 virus has placed extensive strain on RNA isolation and RT-qPCR reagents. Rapid development of new test kits has helped to alleviate these shortages. However, comparisons of these new detection systems are largely lacking. Here, we compare indirect methods that require RNA extraction, and direct RT-qPCR on patient samples. For RNA isolation we compared four different companies (Qiagen, Invitrogen, BGI and Norgen Biotek). For detection we compared two recently developed Taqman-based modules (BGI and Norgen Biotek), a SYBR green-based approach (NEB Luna Universal One-Step Kit) with published and newly-developed primers, and clinical results (Seegene STARMag RNA extraction system and Allplex 2019-nCoV RT-qPCR assay). Most RNA isolation procedures performed similarly, and while all RT-qPCR modules effectively detected purified viral RNA, the BGI system proved most sensitive, generating comparable results to clinical diagnostic data, and identifying samples ranging from 65 copies – 2.1×10^5^ copies of viral Orf1ab/μl. However, the BGI detection system is ∼4x more expensive than other options tested here. With direct RT-qPCR we found that simply adding RNase inhibitor greatly improved sensitivity, without need for any other treatments (e.g. lysis buffers or boiling). The best direct methods were ∼10 fold less sensitive than indirect methods, but reduce sample handling, as well as assay time and cost. These studies will help guide the selection of COVID-19 detection systems and provide a framework for the comparison of additional systems.

## INTRODUCTION

The SARS-CoV-2 coronavirus is a positive-strand RNA virus with a large genome of about 30kb, which encodes up to 14 open reading frames, including several structural genes; e.g. Nucleocapsid (N), Spike (S), Membrane (M) and Envelope (E), accessory genes, and a large open reading frame (Orf1a/Orf1ab) that encodes a polypeptide that is cleaved into 16 non-structural proteins (1, 2). It is related to the SARS-CoV and MERS-CoV coronaviruses, which cause severe respiratory illness in humans, and is the causative agent of the COVID-19 respiratory disease (3). Since the first documented case in Wuhan, China in December 2019, the virus has spread rapidly across the globe. On March 11, 2020, the WHO officially declared COVID-19 a pandemic (4, 5). As of May 12, 2020, there have been over 4.2 million reported cases of COVID-19 and over 286,000 deaths worldwide (6).

The wide range of disease symptoms, including a large portion of mildly or asymptomatic people, has facilitated rapid dissemination (7, 8). Efficient diagnosis, allowing rapid and accurate patient testing remains the key to limiting disease spread. Rapid disease spread has strained the capacity of diagnostic facilities, and the availability of standard reagents. The principle means of diagnostics for COVID-19 relies on RNA extraction from nasal swabs followed by reverse transcriptase-quantitative PCR (RT-qPCR) detection of viral genes (e.g. N, E and RdRp). Rapid development of SARS-CoV-2 RT-qPCR detection systems from many companies has helped to alleviate some of the strain, and many new systems have been given Emergency Use Authorization (EUA) for clinical use. However, comparison of new systems with clinical diagnostics is largely lacking. A limited number of studies have evaluated some kits and compared efficiency of different RT-qPCR primer sets for COVID-19 detection (9–12). These studies have revealed large differences in sensitivity of some systems, highlighting the need for stringent comparison and further optimization of novel detection systems.

An attractive option is direct detection from patient samples without RNA extraction, as it increases throughput, decreases costs and circumvents the need for clinically approved RNA extraction reagents which have become limited. Several studies have examined the ability to directly detect patient samples collected in UTM. While Grant *et al*. reported no effect on sensitivity with extraction-free COVID-19 detection (13), several other studies noted a decrease in the order of 5-20 fold in sensitivity (14–18). Interestingly, while Grant *et al*. observed reduced detection sensitivity after heating the sample to 95°C (13), others have demonstrated that heating samples to 95°C could partially increase sensitivity (14–16), as could detergent-based lysis (16, 19). In studies where large sample numbers were analyzed, optimized extraction-free methods resulted in a high (92-98%) concordance with clinical results, despite reduced sensitivity (14–16).

Here, we comprehensively compare two recently developed COVID-19 detection protocols, one from BGI and the other from Norgen Biotek. The BGI system has been used extensively in China, and has recently been approved for use in several other countries, whereas the Norgen System is currently seeking approval for use starting in Canada. We compare the RNA isolation systems from both companies alongside the Qiagen RNeasy and Invitrogen Purelink systems, both of which are routinely used in research labs, and the former of which has been shown to provide only modestly reduced recovery compared to the CDC approved QIAamp Viral RNA kit (17). We also compare and optimize BGI and Norgen Taqman RT-qPCR detection modules, as well as a SYBR green-based protocol using a commercially available RT-qPCR mix with published and newly designed primer sets. In addition, we evaluate and optimize the ability of the BGI and Norgen systems to detect COVID-19 directly from patient swabs collected in UTM, without RNA extraction. Finally, we perform a cost analysis and discuss both advantages and drawbacks of both systems. We find that both Norgen and BGI RNA isolation kits perform similarly to the Qiagen RNeasy system, while the Invitrogen Purelink is less efficient. We also find that the significantly more expensive, and less flexible BGI RT-qPCR detection module is the most sensitive of the systems tested, providing comparable results to clinical diagnostic data, and could efficiently diagnose patients using extraction-free detection. While initially less sensitive, we found that sensitivity of the more affordable, and flexible Norgen RT-qPCR module could be dramatically improved to levels matching the BGI mix in direct detection assays simply by adding an RNase inhibitor.

## MATERIALS AND METHODS

### Patient samples

Samples were obtained from the MSH/UHN clinical diagnostics lab with approvals from the Research Ethics Boards (REB #20-0078-E) of Mount Sinai Hospital in Toronto, Canada. Clinical diagnostic data was obtained using the Seegene STARMag RNA extraction kit (Microlab STAR Liquid Handling System, Hamilton Company) and Allplex 2019-nCoV RT-qPCR assay analyzed using the Bio-Rad CFX96 IVD real-time qPCR detection system.

### RNA extraction

Qiagen RNeasy, Invitrogen Purelink, Norgen Biotek Total RNA Purification Kit and the BGI Magnetic Bead Viral RNA/DNA extraction kit were used as per manufacturer’s protocols. For each extraction, 100 µl of sample was used and eluted in 32 µl.

### Taqman-based RT-qPCR detection

The 2019-nCoV TaqMan RT-PCR Kit from Norgen Biotek and 2019-nCoV: Real-Time Fluorescent RT-PCR kit from BGI were used essentially as per manufacturer’s instructions. For comparison of different plate formats (Fig. 1A), 10 and 20 µl reaction volumes were used with either 2.5 or 5 µl synthetic RNA standard (Twist Biosci.), respectively, using the Norgen system. These were analyzed in parallel on separate BioRad CFX96 (20 μl reactions) or CFX384 (10 μl reactions) real-time PCR systems. All other experiments used 10 µl reaction volumes (384-well plates) with 2.5 µl of sample (synthetic standard, extracted RNA or direct UTM) and were analyzed using a Bio-Rad CFX384 detection system. For testing alternative primers/probes with the Norgen system, primers/probes were purchased from Integrated DNA Technologies (IDT) and primers were used at 500 nM with probes at 250 nM. Probes were FAM-labelled, E Sarbeco and HKU Orf1 sequences are published (9, 12), and newly designed N gene primers/probe (N Pearson) sequences are Fwd: CCAGAATGGAGAACGCAGT, Rev: TGAGAGCGGTGAACCAAGA, probe: GCGATCAAAACAACGTCGGCCCC). RT-qPCR cycling protocols were as per manufacturers recommendations, except for annealing/elongation temperature testing (Fig. 2E) with the Norgen system where the indicated temperatures were used.

**FIG 1:**
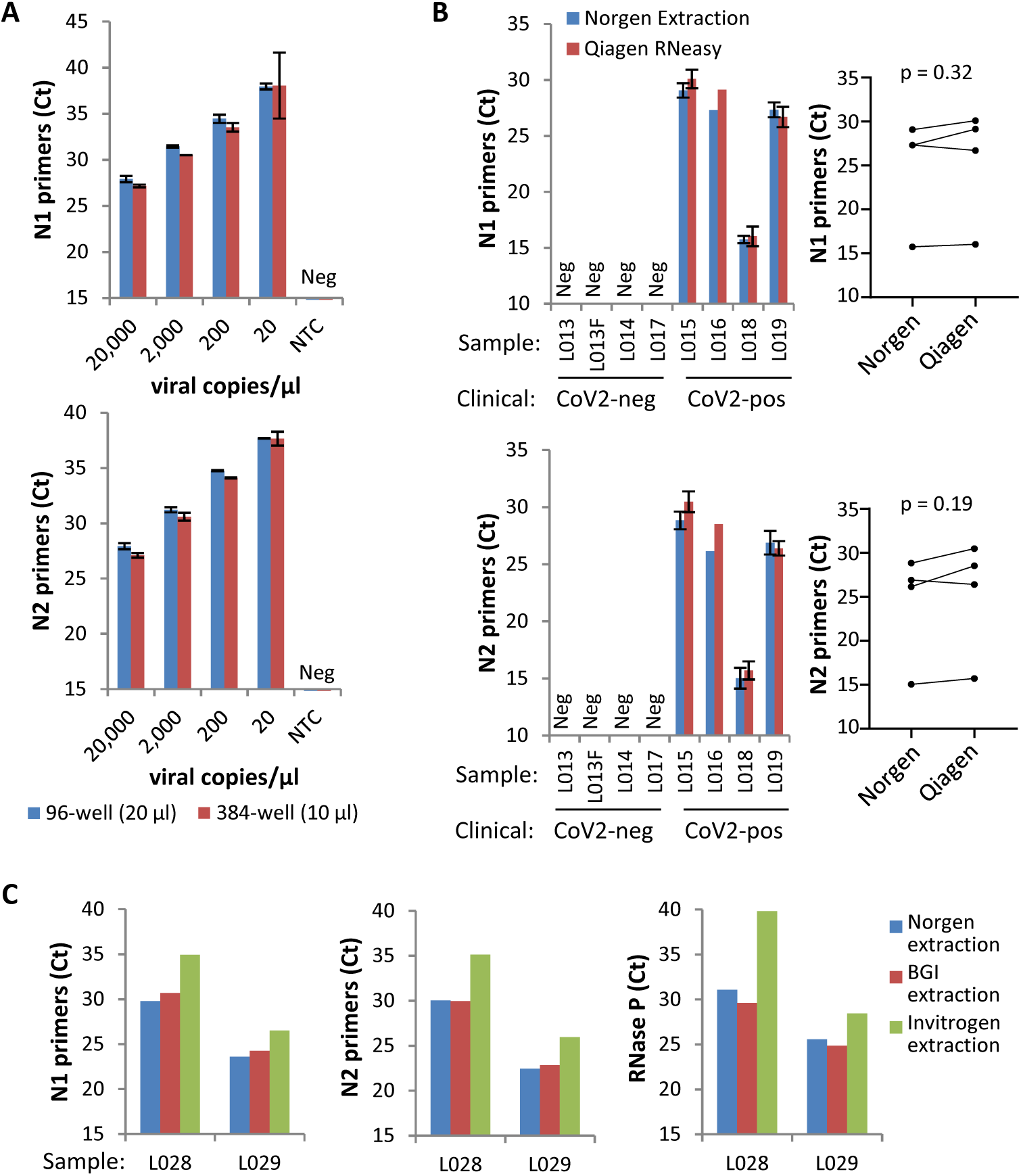
Comparison of plate formats and RNA extraction kits. **(A)** Serial dilutions of SARS-CoV-2 synthetic RNA standards from Twist Biosci (in copies/μl of the standard added to the RT-qPCR reaction) run in parallel on separate BioRad CFX 96-well (20 μl reactions) or 384-well (10 μl reactions) real-time PCR systems using the Norgen COVID-19 RT-qPCR detection module. Mean +/- range of two independent tests. **(B-C)** Analysis of four negative and four positive patient samples extracted with either the Qiagen RNeasy or Norgen RNA isolation kits (B) or two positive patient samples extracted with the Norgen, Invitrogen Purelink or BGI RNA isolation kits (C) using the Norgen RT-qPCR detection system and N1, N2 or human control (Rnase P) primers. Samples L015, L018 and L019 are the mean +/- range of technical duplicates run independently on two separate plates, other samples were analyzed once, although L028 and L029 are rerun in Fig. 2a. In (B) a paired t-test was used to compare Norgen *vs*. Qiagen extractions.

**FIG 2:**
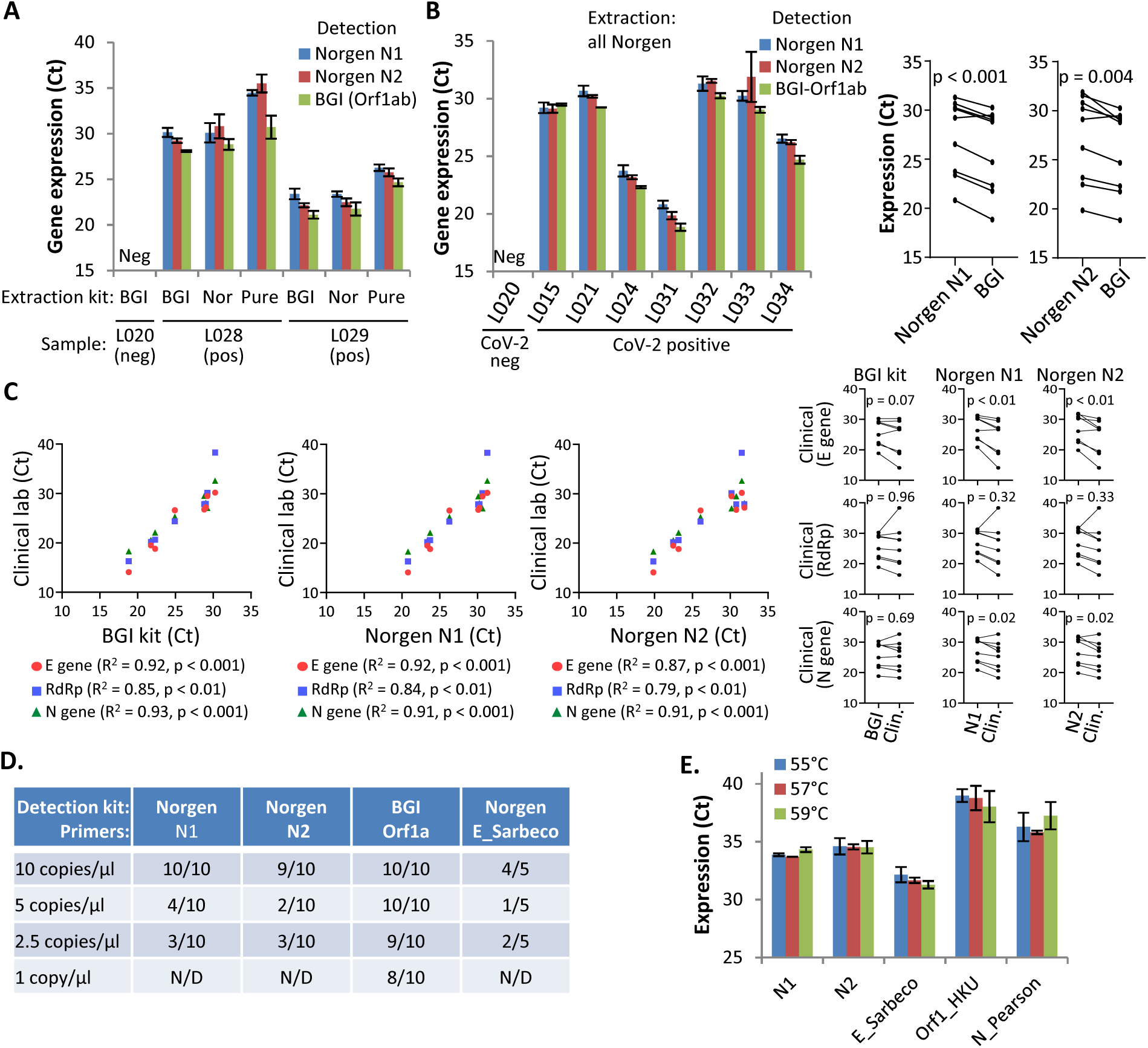
BGI detection kit shows enhanced sensitivity over Norgen kit. **(A)** Analysis of two positive patient samples extracted with the BGI, Norgen (Nor) or Invitrogen Purelink (Pure) RNA isolation kits using the Norgen (N1 or N2 primers) or BGI RT-qPCR detection systems. Mean +/- std dev of the same sample run on three (BGI & Norgen extractions) or two (Purelink extraction) separate plates. **(B)** Analysis of additional patient samples using the Norgen (N1 or N2 primers) or BGI detection systems. Mean +/- range of the same samples run independently on two separate plates. Paired t-tests compare Norgen N1 or N2 and BGI Ct values across all samples (including L028 and L029 extracted with the Norgen kit). **(C)** Comparison of Ct values from clinical lab analysis (E, RdRp and N genes) and data obtained with the BGI or Norgen detection systems. Paired t-tests were used to compare results. **(D)** Detection limit determination using the BGI or Norgen detection systems shown as the number of positives/total number of wells. Concentrations are in copies/µl in the standard. N/D: not determined. **(E)** Analysis of 500 viral copies (Twist Biosci) using N1, N2, E Sarbeco, HKU Orf1 and our N gene (N_Pearson) and the Norgen RT-qPCR mix with the indicated annealing/elongation temperatures. Mean +/- range of two independent tests.

### SYBR green RT-qPCR detection

Primer pairs were designed using PrimerQuest software, and purchased from IDT. Primers selected for testing had ΔG values for self-dimers and heterodimers greater than −9.0 kcal/mole. Newly designed primers were specific for SARS-CoV-2 with no cross-reactivity to other coronaviruses based on published sequences (SH N1 Fwd: AATTGCACAATTTGCCCCCA, Rev: ACCTGTGTAGGTCAACCACG; SH S1 Fwd: TCAGACAAATCGCTCCAGGG, Rev: TCCAAGCTATAACGCAGCCT). The published S gene primers used in this study were S1 Fwd: CCTACTAAATTAAATGATCTCTGCTTTACT, Rev: CAAGCTATAACGCAGCCTGTA (20). Primers were used at 400 nM. RT-qPCR was performed on a LightCycler 480 (Roche) with a 384 well plate using the NEB Luna Universal One-Step RT-qPCR kit (NEB #E3005L, New England Biolabs Inc) and a reaction volume of 10 μl with 2.5 μl of sample. Cycling conditions were as follows: 10 min @ 55°C (RT), 1 min @ 95°C (denaturation), 45 cycles: 10s @ 95°C, 30s @ 60°C (amplification), melt curve. Standard curves were generated for each primer set with serial dilutions of viral RNA from 0.8 to 800,000 copies/μl; SARS-CoV-2 RNA (strain USA_WA1/2020) was provided by the World Reference Centre for Emerging Viruses and Arboviruses (Galveston, TX) (WRCEVA).

### Direct extraction-free SARS-CoV-2 detection

For direct detection, 2.5 μl of patient sample in universal transport media (UTM, Copan) were added to the RT-qPCR reaction mix. For comparison to extracted RNA, an equivalent input of extracted RNA was used (i.e. extracted RNA eluted in 32 μl was diluted 1:2 with RNase-free water). To optimize direct detection, RNaseOUT (ThermoFisher) was added to UTM samples (2 U/μl). Samples were then left untreated, heated at 95°C for 15 min, mixed 1:1 with Lucigen QuickExtract DNA extraction solution with heating at 95°C for 5 min or treated with MyPOLS Bio VolcanoCell2G lysis buffer, 1% Triton X-100, 1% Tween-20 or 1% Saponin and incubated on ice 15min. Samples were then directed added to the RT-qPCR reaction mixture and compared to UTM samples that had been left untreated.

## RESULTS

### Plate format

Many diagnostic protocols utilize 20 µl reactions in 96-well plates, but reducing volume in a 384-well format increases throughput and reduces costs. Using the Norgen RT-qPCR COVID-19 detection kit (which utilizes CDC-approved N1 and N2 primers), we observed similar Ct values in a comparison of 20 vs 10 µl reactions in 96- or 384-well plates, respectively (Fig. 1A), thus in subsequent analysis we focused on 384-well plates.

### RNA Extraction Methods

Qiagen RNA extraction systems are used extensively for viral RNA isolation, but availability has become limited. Thus, we first compared the Qiagen RNeasy RNA extraction kit to a similar kit from Norgen Biotek, both of which utilize silica-based columns. None of the SARS-CoV-2-negative samples generated any signal, and we detected no significant difference in Ct values across four clinically-diagnosed positive patient samples (Fig. 1B), thus the Norgen extraction system performs similarly to standard Qiagen kits. We next compared efficiency of the Norgen (column based), Invitrogen Purelink (column-based) and BGI (magnetic bead-based) RNA isolation systems. Using two new positive patient samples, we observed similar recovery with both the Norgen and BGI systems, but considerably higher Ct values were observed for viral (N1 and N2 primers) and human control (RNase P) genes with the Invitrogen kit (Fig. 1C). Thus, for isolating SARS-CoV-2 RNA from nasopharyngeal patient samples in UTM, Norgen, Qiagen RNeasy and BGI extraction methods are all comparable, but the Invitrogen Purelink kit is less efficient.

### TaqMan Primers/RT-qPCR mix

Next we compared the efficiency of two recently developed TaqMan-based SARS-CoV-2 RT-qPCR detection kits from Norgen and BGI. The BGI protocol uses one primer set against Orf1ab, while as noted above, Norgen uses two separate reactions targeting the N gene. We first compared RNA from the two positive patient samples extracted with the Norgen, BGI or Invitrogen Purelink methods, as well as RNA isolated from one negative patient sample using the BGI extraction method. These assays confirmed that Norgen/BGI extraction methods are more efficient than Purelink, but also revealed greater sensitivity (∼1-3 Ct values) with BGI *vs* Norgen primers/RT-qPCR mix, particularly with the lowest level sample (L028-Purelink) (Fig. 2A). We observed a similar trend across seven additional samples all isolated using the Norgen RNA extraction kit (Fig. 2B). Comparison to clinical values for the viral N, E and RdRp genes obtained using the Seegene STARMag RNA extraction kit and Allplex 2019-nCoV RT-qPCR assay analyzed using the Bio-Rad CFX96 IVD real-time qPCR detection system, showed a strong correlation with Ct values obtained using either the BGI or Norgen detection modules (Fig. 2C). We observed no significant difference between the BGI and clinical values, although there was a trend toward lower Ct values with the clinical lab E gene primers/probe, particularly with higher-level samples (Fig. 2C). In contrast, the Norgen detection module showed significantly higher Ct values compared to clinical data (median 1.4 to 3.5 Ct values higher depending on the primer sets used) (Fig. 2C), similar to what we observed in comparison to the BGI detection system. Using synthetic TWIST Bioscience SARS-CoV-2 standards we found that the BGI detection kit routinely detected 2.5 copies/µl (6.25 copies/10µl rxn), and 1 copy/µl (2.5 copies/10µl rxn) 80% of the time, whereas the detection limit of the Norgen system was 10 copies/µl (25 copies/10µl rxn) (Fig. 2D), with lower concentrations being detected < 50% of the time.

To determine if sensitivity using the Norgen RT-qPCR mix could be enhanced, we tested different annealing/elongation temperatures in the qPCR reaction along with two other published SARS-CoV-2 primers/probes shown to have high sensitivity (E Sarbeco and HKU Orf1) (9, 12, 21, 22), and new primers/probes we designed to target the viral N gene. Increasing the annealing/elongation temperature from the manufacturer’s recommended 55°C did not affect Ct values for either the N1 or N2 primers provided with the Norgen system (Fig. 2E). Using the Norgen RT-qPCR mix, we observed poor performance of the HKU Orf1 primer set, and the newly designed N gene primers provided higher Ct values compared to the CDC N1 and N2 primers, but the E gene primers/probes demonstrated lower (more sensitive) Ct values compared to the N1/N2 primers, particularly at 59°C annealing/elongation (Fig. 2E). This improvement, however, did not translate to a lower detection limit (Fig. 2D). Thus, while both systems easily detect purified SARS-CoV-2 RNA from infected patients, the BGI primers/RT-qPCR system provides a lower detection limit and similar Ct values to clinical data, while Ct values for the Norgen detection module are ∼2-3 cycles higher.

### SYBR green detection

We next compared the more sensitive BGI detection system to a SYBR green-based method. We tested various published primers, some designed for SYBR green and some from TaqMan assays (9, 12, 20, 23), and designed our own. One published set for the viral S gene (20) and two new N or S gene primer sets gave little/no signal in no-template control (NTC) and generated a linear response across 8 - 800,000 viral copies/μl (Fig. S1A), and were thus selected for future analyses. We then compared SARS-CoV-2 standards using the SYBR green primers and the BGI detection kit and observed comparable Ct values between the two systems across 20 to 20,000 genome copies/μl (Fig. 3A). Identical Ct values were obtained using SARS-CoV-2 RNA from WRCEVA (not shown). The BGI system provided a slightly lower detection limit than the SYBR green systems (compare Figs. 2D and 3D).

**FIG 3.**
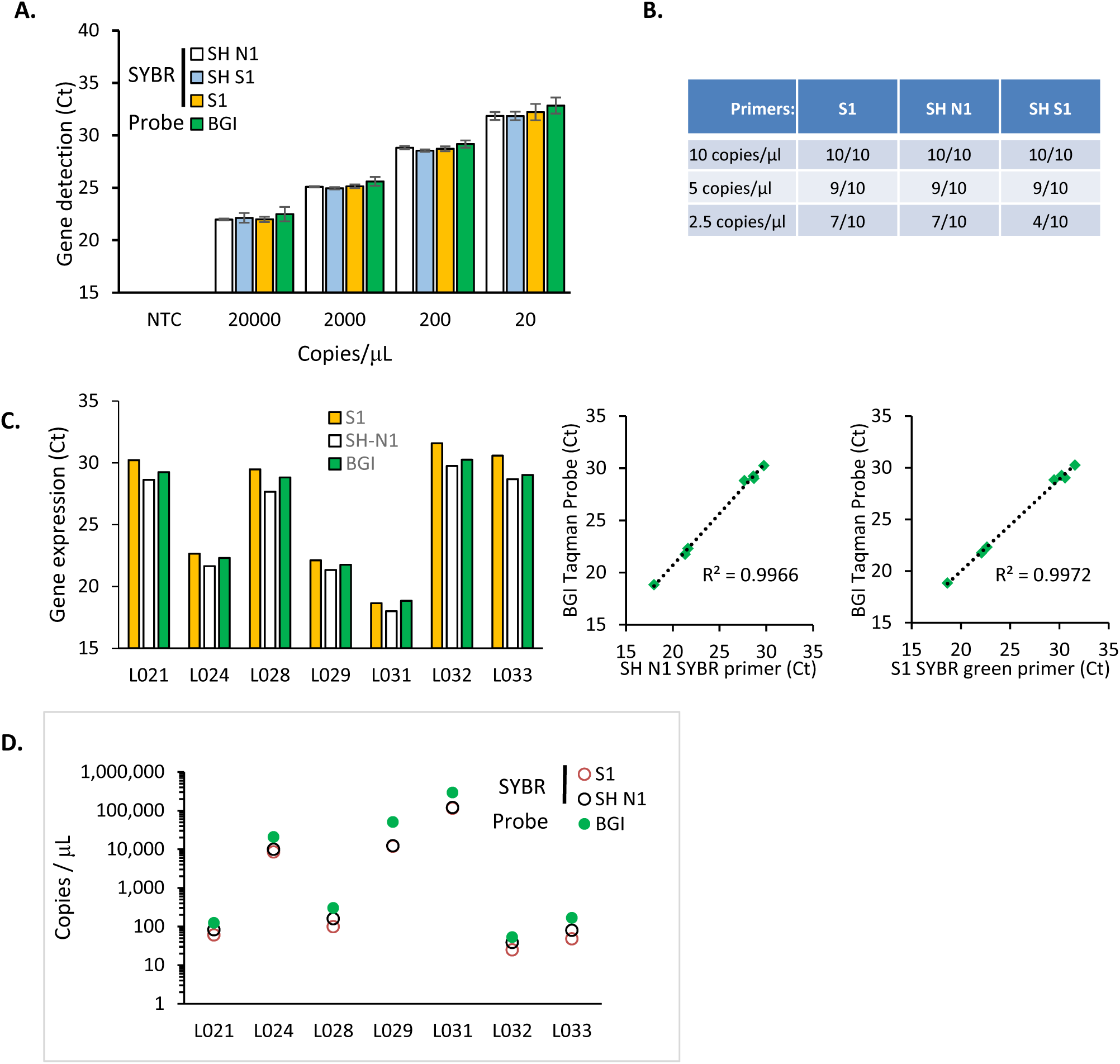
SYBR green detection of extracted RNA. **(A)** Serial dilutions of SARS-CoV-2 synthetic RNA standards (Twist Biosci) were run in SYBR green and BGI TaqMan assays. Mean +/- STD; n ≥ 3. **(B)** Detection limit for each of the SYBR green primer sets shown as the number of positive samples/total number of samples tested. Synthetic RNA (Twist Biosci) was used from stocks with the indicated number of copies per μL. **(C)** Comparison of Ct values obtained for each patient sample with the SYBR green and BGI TaqMan assays. Linear regression was used to determine the R2. BGI data is from Fig. 2A, B. **(D)** Comparison of viral copy number per μL for each of the positive patient samples determined with each primer set. Copy number was determined using a standard curve of SARS-CoV-2 RNA (WRCEVA).

We next analyzed patient samples comparing the SYBR green primers to previous data obtained with the BGI kit (Fig. 3C). One of the primers (SH S1) did not perform well on patient samples and was excluded from these experiments. The other SYBR green primers reliably identified all negative and positive patient samples, with SH-N1 primers generating slightly lower Ct values (0.3 to 1.1 Ct values, p = 0.02) and S1 primers providing slightly higher Ct values compared to the BGI system (−0.2 to 1.6 Ct values, p < 0.01). Quantification of gene copy numbers generated similar results for SYBR green and BGI, and ranged from 24 copies to >120,000 copies/µl (Fig. 3D). Non-specific melt peaks were occasionally observed in negative and low virus copy positive samples, which could easily be identified and excluded (Fig. S1B). All patient samples were positive for human RNase P (not shown).

### One step detection without RNA purification

To reduce the number of steps required for viral detection we tested RT-qPCR direct from patient samples in UTM. For this, we added 2.5 μl of sample directly to the RT-qPCR mix and compared this to an equivalent input of extracted RNA. UTM blocked SYBR-green detection of SARS-CoV-2 RNA standards (data not shown), but both the BGI and Norgen TaqMan detection systems identified positive patient samples (Fig. 4A). Ct values were lower for BGI *vs* Norgen, consistent with data with purified RNA (*c*.*f*. Figs. 2 & 4A). Furthermore, the Norgen system did not reliably identify some positive samples with lower levels of virus (Fig. 4A). Relative to extracted RNA, direct RT-qPCR with the BGI detection kit was 2-26 fold less sensitive (except sample L021, which was ∼600-fold reduced, see below for an explanation), whereas with the Norgen kit it was 20-1000’s fold lower (L033 with the N2 primers was an exception at 4.4-fold). Despite the reduced sensitivity, the strong correlation between BGI and clinical Ct values was maintained (Fig. 4B).

**FIG 4:**
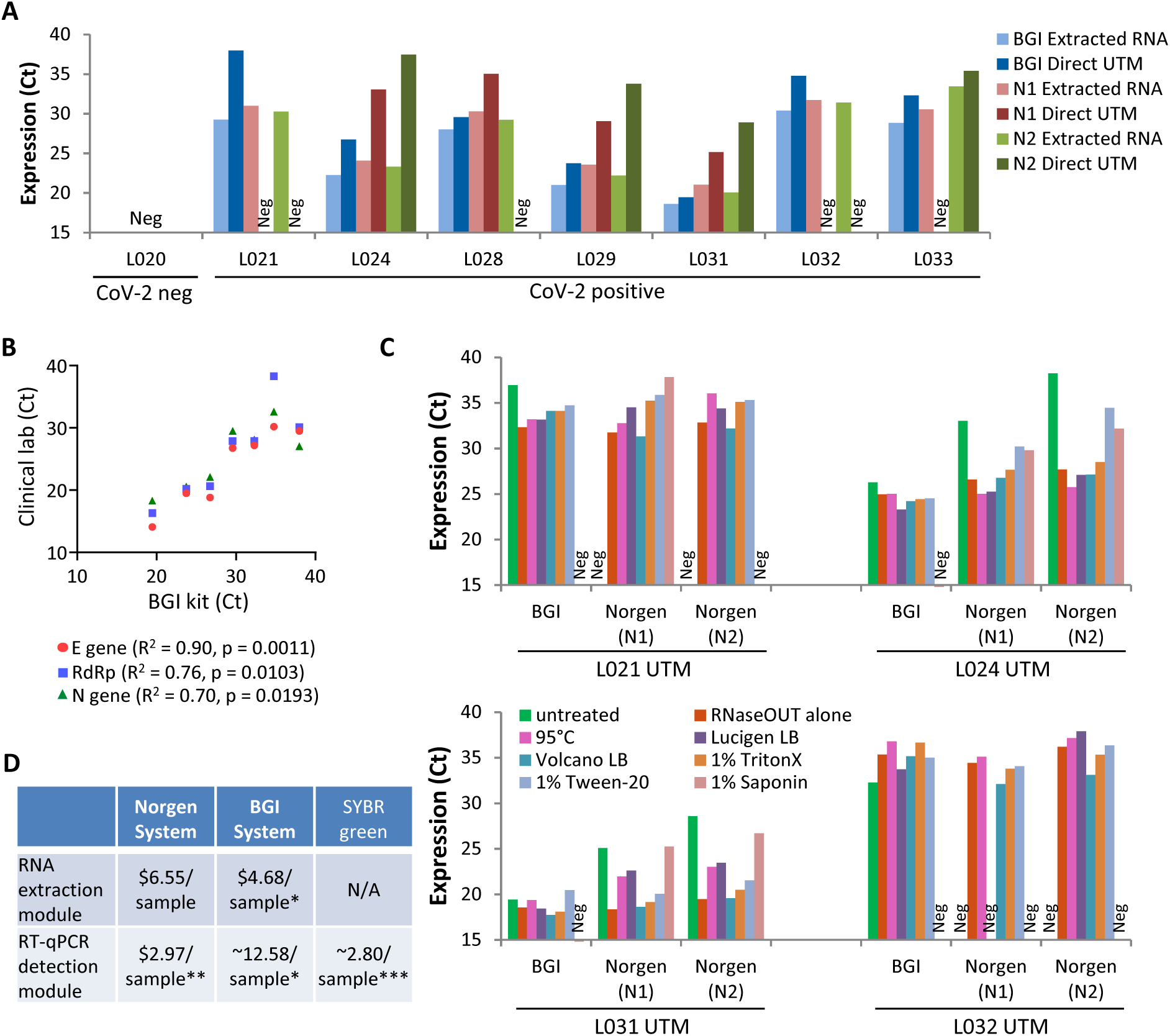
One-step direct detection without RNA extraction. **(A)** Analysis of extracted RNA or direct UTM from a panel of patients using the BGI or Norgen (N1 and N2 primers) detection systems. **(B)** Comparison of Ct values from clinical lab analysis on extracted RNA (E, RdRp and N genes) to data obtained for direct analysis with the BGI detection system. **(C)** Patient samples in UTM were left untreated, or treated with the RNase inhibitor RNaseOUT with or without heating at 95°C for 15 min, or treated with the indicated lysis buffers/detergents and then directly analyzed using the BGI or Norgen (N1/N2 primers) RT-qPCR detection systems. Note sample L020 (clinical negative) was also tested under these conditions and was confirmed as SARS-CoV-2 negative. **(D)** Cost analysis comparing Norgen, BGI, and SYBR green systems. Price is in CAD at the time these studies were initiated (late March/early April 2020) for 10µl RT-qPCR reactions and include relevant processing and shipping fees. * BGI RNA extraction module is based on the 96-sample format, price can be reduced ∼15% by purchasing the 1728-sample format, and bulk pricing with a ∼25% discount of the detection module is available for >10,000 samples. ** Pricing for the Norgen detection module is based on the 50-sample format running three separate wells (N1, N2 and RNaseP) per sample, pricing can be reduced if purchasing the larger 500-sample format. *** Pricing for SYBR green detection is based on the 200 reaction size LUNA Universal One-Step RT-qPCR Kit (NEB) running three separate wells/sample (two viral genes and one human control gene). Pricing can be reduced up to 30% with larger kit sizes. N/A: not applicable.

Others have reported that reduced sensitivity in direct *vs*. extracted RNA analyses can be partially overcome by heat or different lysis buffers/detergents (14–16, 19). Thus, we assessed the effect of adding an RNase inhibitor (RNaseOUT), heating samples at 95°C for 15 minutes, or five different lysis buffers/detergents (Lucigen QuickExtract DNA extraction solution, MyPOLS Bio VolcanoCell2G lysis buffer, 1% Triton X-100, 1% Tween-20 or 1% Saponin). Simply adding RNase inhibitor was sufficient to dramatically increase detection >100 fold using the Norgen system, bringing Ct values to levels comparable to those obtained with the BGI RT-qPCR system, and, most importantly, allowing for detection of previous “false-negative” samples L021 and L032 (Fig. 4C). Furthermore, RNase inhibitor brought direct RT-qPCR results with the Norgen detection kit to within 3 Ct values (∼10-fold) of those obtained with extracted RNA (compare Figs. 4A and C). Treatment with heat, lysis buffers or detergents did not appreciably increase sensitivity further, and in some cases reduced sensitivity (higher Ct values). For the BGI detection system, none of the treatments dramatically improved detection, with the exception of sample L021 (Fig. 4C), which previously showed the largest difference between extracted RNA and direct UTM analysis (Fig. 4A). We presume, therefore, that L021 had higher RNase levels that were not fully inhibited by the (proprietary) RNase inhibitor already present in the BGI mix. Thus, RNase inhibitor is sufficient to improve direct detection and under these conditions BGI and Norgen kits perform similarly.

## DISCUSSION

Here, we comprehensively compared four different RNA isolation methods, two recently released SARS-CoV-2 TaqMan RT-qPCR detection modules and a SYBR green-based RT-qPCR approach for SARS-CoV-2 detection using published and newly-developed primers. Furthermore, we tested and optimized extraction-free SARS-CoV-2 detection using these same detection modules.

For RNA extraction, we tested three different column-based systems from Qiagen (RNeasy), Invitrogen (Purelink) and Norgen Biotek, as well as a magnetic silica bead system from BGI. While only the BGI system is specifically marketed for viral RNA isolation, we observed similar results using the Qiagen RNeasy, Norgen and BGI systems, and while it was only tested on two samples, we observed lower recovery of viral RNA using the Invitrogen Purelink system. Cost analysis of the BGI and Norgen Biotek RNA isolation systems revealed that the latter is ∼40% more expensive than that of BGI ($6.55 CAD *vs*. $4.68 CAD/sample, Fig. 4D), but we found that for small batches of samples the bead-based BGI kit was slower, increasing sample preparation time by about 50% over the Norgen kit (∼30 *vs*. 45 min). This difference was largely due to two incubation steps in the BGI protocol, so the relative difference in sample preparation time may diminish as larger numbers of samples are processed. Furthermore, magnetic beads facilitates large-scale, automated sample extraction.

For RNA detection, we tested TaqMan-based detection systems from BGI and Norgen Biotek, as well as a SYBR green method using a commercially available RT-qPCR mix and published primers (some used for SYBR green and others from probe-based methods) along with new primers we developed. All systems could accurately detect SARS-CoV-2 positive patient samples using extracted RNA, and generated Ct values that strongly correlated with clinical diagnostic values. However, the BGI and SYBR green methods routinely produced lower Ct values for patient samples, which closely match clinical results, and had lower detection limits compared to the Norgen system. The BGI system also performed slightly better that the SYBR green methods with low-level standards. One drawback to the SYBR green method was reduced specificity, as we sometimes observed non-specific products in negative or low-level samples, although these could be identified by monitoring melt curves. These non-specific products were not routinely observed in NTC reactions. Thus, melt curve analysis is an essential component of SYBR green qPCR. We also tested 8 other published and newly designed primers and all yielded non-specific PCR products (not shown). Whether non-specific products can be eliminated using alternative RT-qPCR mixes remains to be determined. The BGI detection module is over four-times more expensive than Norgen or SYBR green methods (Fig. 4D), providing a significant financial drawback. Cost savings with the Norgen kit could be even greater if multiplexing primers/probes were utilized; currently this system follows the CDC guidelines with three separate reactions, one each using FAM-labelled viral N1, viral N2 or human RNase P primers/probes. The Norgen and SYBR green systems also provide more flexibility than that of BGI. Primers/probes come pre-mixed in the BGI system and cannot be altered, whereas they are added separately in the others, allowing alternative primer/probe options and concentrations. We tested three alternative primers/probes with the Norgen system. Those targeting the E gene performed similarly to the provided N1/N2 primers/probes, while alternatives for the viral N or Orf1a gene had reduced sensitivity, although only a single primer/probe concentration was tested. Sequences of the BGI primers/probes are unavailable, and only a single primer/probe set targeting the viral Orf1ab gene is used. Mutation could affect detection and generate false negatives. Thus, while the BGI system provides a lower detection limit with extracted RNA than Norgen or SYBR green detection systems, all accurately identified SARS-CoV-2-positive patients, and latter systems detect multiple viral targets and offer greater flexibility and substantially reduced costs.

Finally, we tested direct, extraction-free detection of SARS-CoV-2. This approach reduces cost, increases throughput, and circumvents the need for RNA extraction systems that may be scarce during a pandemic. Others have shown that SARS-CoV-2 can be detected from patient samples, although this typically comes with reduced sensitivity, which can at least partially be overcome by heat and/or detergent lysis (14–16, 19). We found that SYBR green-based detection was incompatible with direct detection of samples in UTM. The unmodified BGI detection system performed well in the direct detection of unprocessed patient samples, and confirmed all positive samples tested across a wide range of clinical values, but had a reduced median sensitivity of ∼12-fold compared to extracted RNA. The Norgen system initially performed poorly on direct UTM samples, generating much higher Ct values than extracted RNA (in some cases 1000s of fold higher), and resulted in several false-negatives. Critically, however, adding RNase inhibitor increased sensitivity of direct RT-qPCR with the Norgen system > 100-fold, allowing detection of all previously false-negative samples. This modification did not, in most cases, dramatically increase sensitivity of direct sample analysis with the BGI detection system, suggesting it already contains an RNase inhibitor. Even in that case however, detection of one patient sample was markedly improved, implying higher RNase levels. Thus, addition of RNase inhibitor is a simple and sufficient step to facilitate diagnosis of SARS-CoV-2 direct from patient samples.

Our results provide in depth analysis of recently released SARS-CoV-2 detection systems from BGI and Norgen Biotek and compare these to a SYBR green-based approach and to clinical diagnostic values. Each system provides advantages and disadvantages depending on sensitivity, specificity, flexibility and cost. Our findings will help guide selection of SARS-CoV-2 detection systems, and provide an outline for others to compare alternative systems.

## ACKNOWLEDGEMENTS

This work was funded by a grant to RB, JW and LP from the Krembil Foundation, and the Canadian Institutes of Health Research (CIHR; FDN-154299) to MIC.

## FIGURE LEGENDS

**Supplementary Figure S1:**
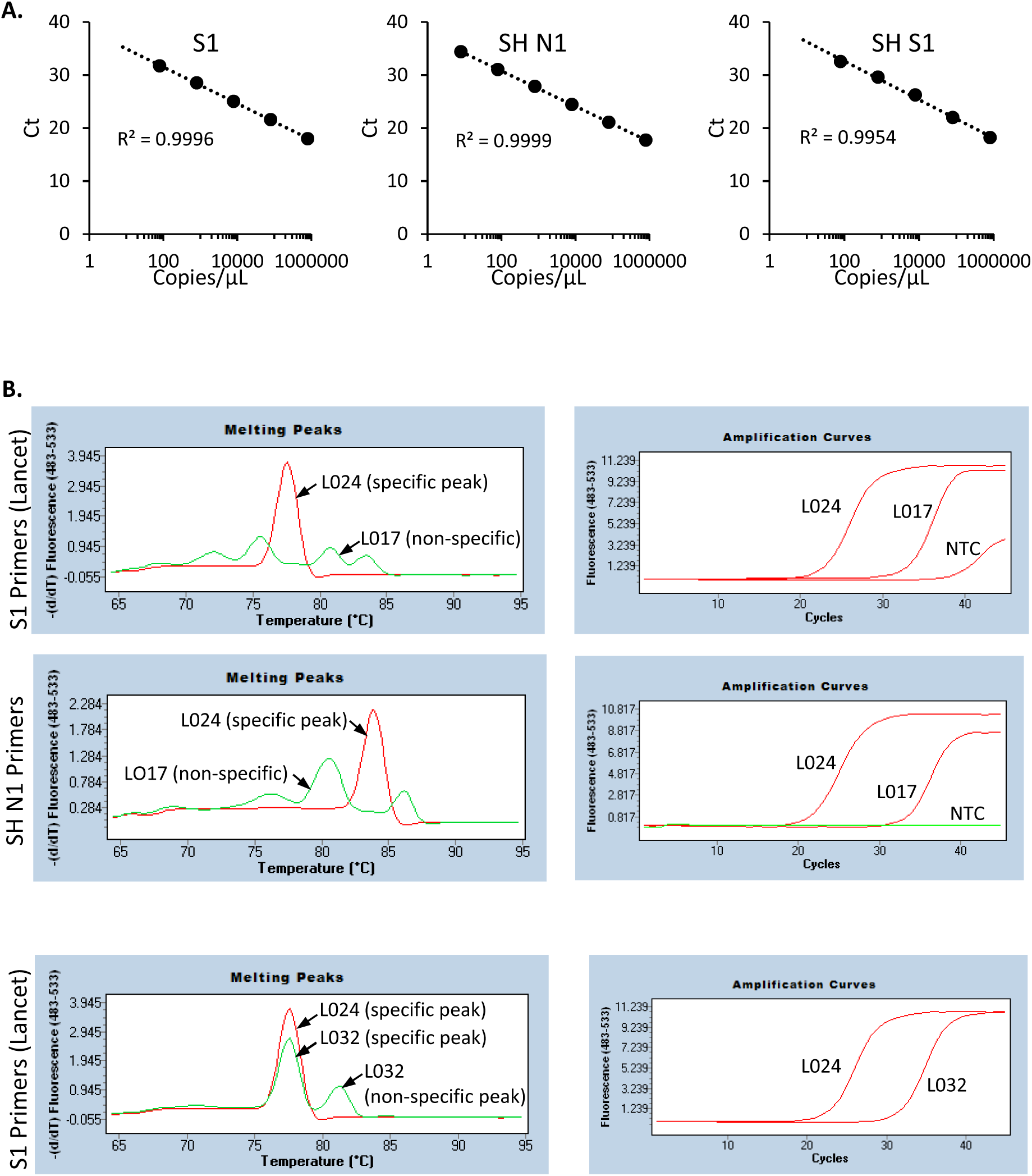
SYBR green detection of SARS-CoV-2. **(A)** Standard curves were generated with 8 to 800,000 copies of SARS-CoV-2 RNA (WRCEVA). Mean, n = 5, 2 independent experiments. **(B)** Examples of melt curves from a positive high SARS-CoV-2 copy number sample (L024) showing a single specific melt peak, a negative sample (L017) showing non-specific melt peaks, and a positive low SARS-CoV-2 copy number sample (L032) showing both specific and non-specific melt peaks. NTC, no template control (water).

